# CDK5 influences the organization of the Circadian Machinery in peripheral clocks

**DOI:** 10.1101/2021.10.01.462714

**Authors:** Jürgen Ripperger, Urs Albrecht, Andrea Brenna

## Abstract

Circadian rhythms are self-sustained physiological changes that drive rhythmicity within the 24-hours cycles. Posttranslational modifications (PMTs), such as protein phosphorylation, acetylation, sumoylation, and ubiquitination, are biochemical processes that modify protein structure and functions, ensuring circadian rhythm precision. For example, phosphorylation is considered the most important hallmark of rhythmicity from cyanobacteria to mammals. Cyclin-dependent kinase 5 (CDK5) has been shown to regulate the mammalian SCN’s circadian clock via phosphorylation of PER2. Here, we show that CDK5 influences the clock machinery assembling, using immortalized mouse embryonic fibroblast as an in *vitro* model for studying the peripheral clock. In fact, the circadian period at the cellular level is lengthened. Furthermore, the clock-controlled gene’s expression amplitude is dampened in *Cdk5* ko cell lines, while the phase is delayed about 4 hours.

Taken together, we show *in vitro* that CDK5 is critically involved in regulating the peripheral clocks, influencing their temporal and spatial dynamics.

## Introduction

Circadian rhythms consist of interconnected transcriptional/translational and posttranslational feedback loops that dictate the rhythmicity of physiological responses within 24-hours (Schibler and Sassone-Corsi, 2002; Brenna and Albrecht 2020). These rhythms can be synchronized by stimuli called zeitgebers (ZT). For instance, in mammals, the light can reset the rhythmicity in the suprachiasmatic nuclei (SCN), a bipartite brain region belonging to the hypothalamus, placed above the optic chiasm. Thus, the SCN maintains the internal rhythms synchronously with the dark-light cycles. In addition, the SCN coordinates the synchronization of the independent circadian oscillators situated in peripheral tissues (i.e., liver, kidney, heart) through nervous and hormonal signaling pathways, regulating a wide range of physiological responses (Dibner et al., 2010). The rhythmic transcription is driven by the core factors of the positive loop, the heterodimer CLOCK: BMAL1. This heterodimer drives the expression of many clock-controlled genes (CCGs), such as *Periods* (Per 1, 2, 3) and *Cryptochromes* (Cry 1, 2) by binding the E-box element (CACGTG) within their promoter (Partch et al., 2014). On the other hand, PERs and CRYs protein products are involved in the negative feedback loop. After protein translation, PER: CRY homo/heterodimers translocate into the nucleus and inhibit CLOCK: BMAL1–mediated transcription through direct protein-protein interaction (Buhr and Takahashi, 2013). Thereby, the cellular positive and negative feedback loop promotes the oscillation of about 10–20% of all genes expressed (Panda et al., 2002a; Storch et al., 2002). Additionally, the fine-tuned period of the circadian oscillators is ensured by many additional levels of regulation, such as epigenetic and posttranslational modifications (PMTs) (Bellet and Sassone-Corsi 2010; Hirano et al., 2016). Within the PMTs, phosphorylation is the most important hallmark of rhythmicity (Roblest et al., 2017). In some organisms, such as cyanobacteria, the circadian rhythmicity is driven only by phosphorylation, without the addition of transcriptional/translational loops (Iwasaki et al., 2002).

In mammals, many kinases are involved in regulating circadian rhythms (Brenna and Albrecht, 2020). Cyclin Dependent-Kinase 5 is a serine/threonine kinase belonging to the Cdc2/Cdk1 family activated by specific cofactors such as p35 and p39 (Tang et al., 1995; Tsai et al., 1994) and cyclin I (Brinkkoetter et al.,2009). CDK5 plays many roles in the brain, such as neurogenesis, neuronal migration, axon guidance, and aberrant activity leads to many neurodegenerative diseases (Kawauchi, 2014). CDK5 can phosphorylate both CLOCK and BMAL1 (Kwak et al., 2013; Brenna et al., 2019). This kinase influences in the SCN the PER2 protein stability and interaction with CRY1, which are the core elements of the negative loop, ensuring the gene expression rhythmicity. Lack of CDK5 shortens the circadian period length in mice (Brenna et al.,2019). Therefore, this kinase has been proposed as one of the main regulators of circadian rhythmicity in mice (Brenna and Albrecht 2020). In addition, Although it was identified for the first time as one o the main kinases playing a role in the brain, many recent observations led to the discovery of the involvement of CDK5 pathways outside the brain. CDK5 regulates vesicular transport, apoptosis, cell adhesion, and migration ((Contreras-Vallejos et al., 2012). Dysfunction of CDK5 in peripheral tissues is associated with many forms of cancer also connected to the aberration of the circadian clock (Pozo and Bibb 2016; Sulli et al., 2019).

Here we show for the first time the involvement of CDK5 in the regulations of the peripheral clock, using immortalized mice embryonic fibroblasts as a model. Our results show that the kinase activity of CDK5 is circadian, and deletion of this enzyme leads to a prolonged period length. Furthermore, this phenotype is associated with an aberrant shuttling into the nucleus of the clock factors. Thus, the BMAL recruitment to the E-box is shifted, and gene expression is altered. Altogether our results show that CDK5 is essential for the correct assembly of the clock machinery.

## Results

### The kinase activity of CDK5 in immortalized mouse embryonic fibroblasts is circadian

CDK5 kinase activity in the SCN is diurnal, with an off/on the state under light: dark (12:12) conditions (Brenna et al.,2019). To understand whether the enzymatic activity can be circadian, we challenged the hypothesis using the NIH3T3 (immortalized mouse embryonic fibroblasts) cell lines as a model system. We synchronized cells with forskolin, a powerful zeitgeber for circadian rhythmicity (Yagita and Okamura 2000), and collected samples at different circadian time (CT) points. We validated the circadian entrainment of cells by immunoblotting, using antibodies against CRY1 and BMAL1 (Fig. 1A). CDK5 was subsequently immunoprecipitated at each time point, followed by *in vitro* kinase assay using histone H1 as substrate in the presence of radiolabeled ATP. The autoradiography signal showed that CDK5 kinase activity is circadian with an “on” state at CT 12-16 and 28-32 (Fig. 1B, C). These CTs correspond to the subjective nights. The specificity of the reaction was verified by repeating the assay by immunoprecipitating the kinase from *Cdk5 KO* cell lines or wt cells pretreated with Roscovitine 34 uM (a CDKs inhibitor) (Tomov et al., 2019), followed by *in vitro* kinase assay. The phosphorylation of H1 did occur only in the wt samples (Fig. 1D).

**Figure 1.**
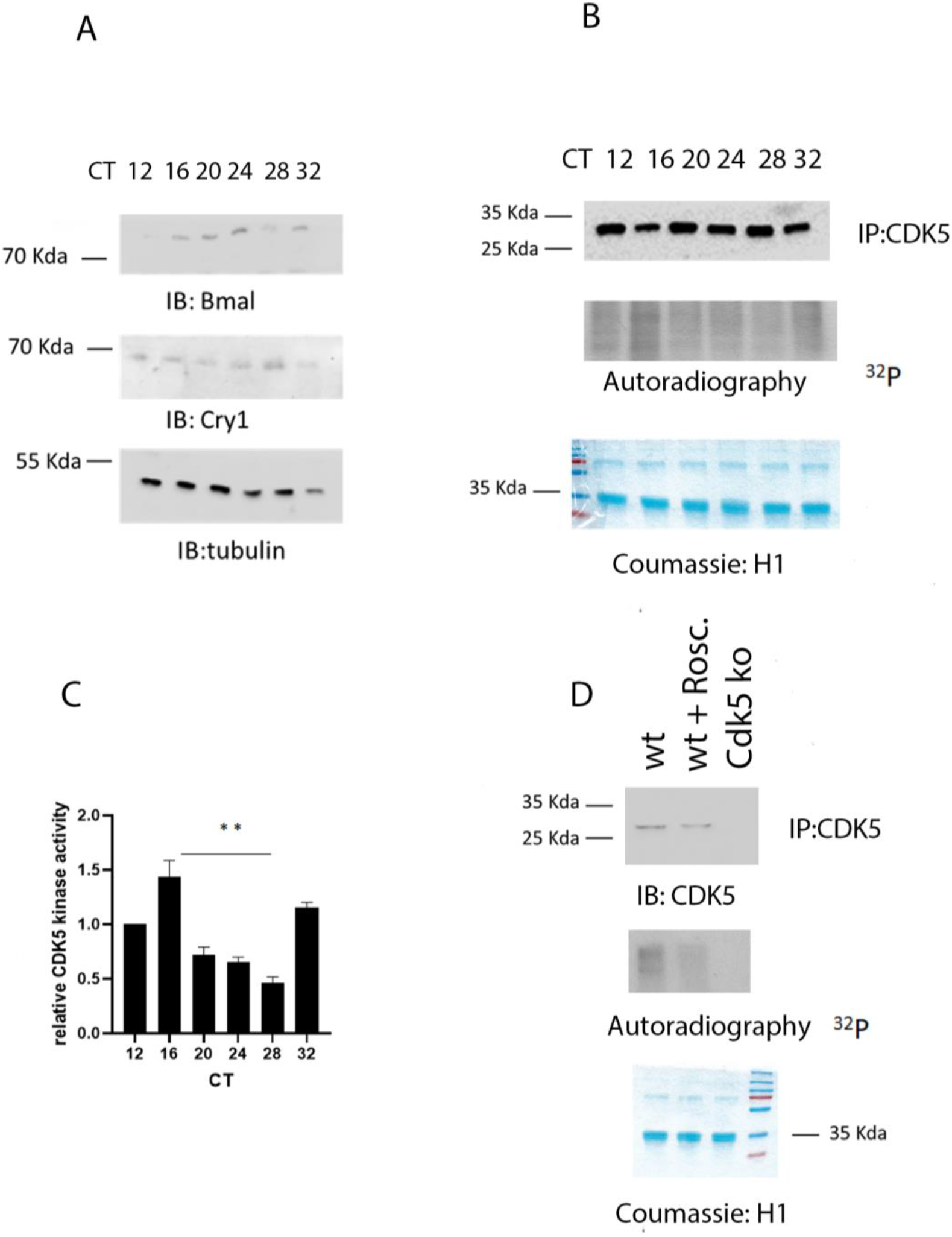
CDK5 kinase activity is circadian. A) Western Blot performed on forskolin-synchronized wt cell, collected at different circadian points (CTs). Samples were stained with antibodies against CRY1 and BMAL1 to validate the circadian cellular synchronization. B) CDK5 was immunoprecipitated at different CTs, and subsequently, an *in vitro* kinase assay was performed using histone H1 as substrate in the presence of radiolabelled ^32^P^-^ATP. C) The radioactive signal was normalized on the immunoblotted signal for CDK5 after the IP. Quantification of three independent experiments (mean ± SEM). One-way ANOVA with Bonferroni’s post-test, *: p<0.001. D) the experiment in (B) was performed in the presence of cells previously treated with Roscovitine and a *Cdk5* ko cell line. The experiment confirmed the specificity of the phosphorylated signal due to the CDK5 kinase activity.

Altogether these observations support the thesis that CDK5 phosphorylates the histone H1 in a circadian fashion.

### Lack of CDK5 lengthens period in peripheral clock

To understand whether CDK5 can affect the circadian period in the peripheral clock, we transfected NIH3T3 with *Bmal*^*luc*^ promoter and treated cells either with DMSO (vehicle) or Roscovitine (CDKs inhibitor) at different concentrations for 12 hours. This treatment was followed by synchronizing the cells with forskolin and recording the real-time luciferase bioluminescence over days (Fig 2A). NIH3T3 cell lines treated with DMS0 show a regular period length, which settles around 23 hours while increasing amount of the drug corresponded to an increased period length (Fig 2B). To confirm the involvement of CDK5, we subsequently modulated the Cdk5 gene expression transfecting the *Bmal*^*luc*^ promoter in NIH3T3 cell lines with different sh RNA against *Cdk5* mRNA (Brenna et al., 2019 Fig. S1A). The increasing effect of the sh RNA against *Cdk5* mRNA mirrored the lengthening of the circadian period in cells (Fig 2C, D). Thus, we finally confirmed that lack of CDK5 lengthens the period, comparing NIH3T3 vs. *Cdk5* KO cell lines (Fig. 2E, 2F). Altogether, these results suggest that lack of CDK5 has a tissue-specific effect in the circadian period of the peripheral clocks

**Figure 2.**
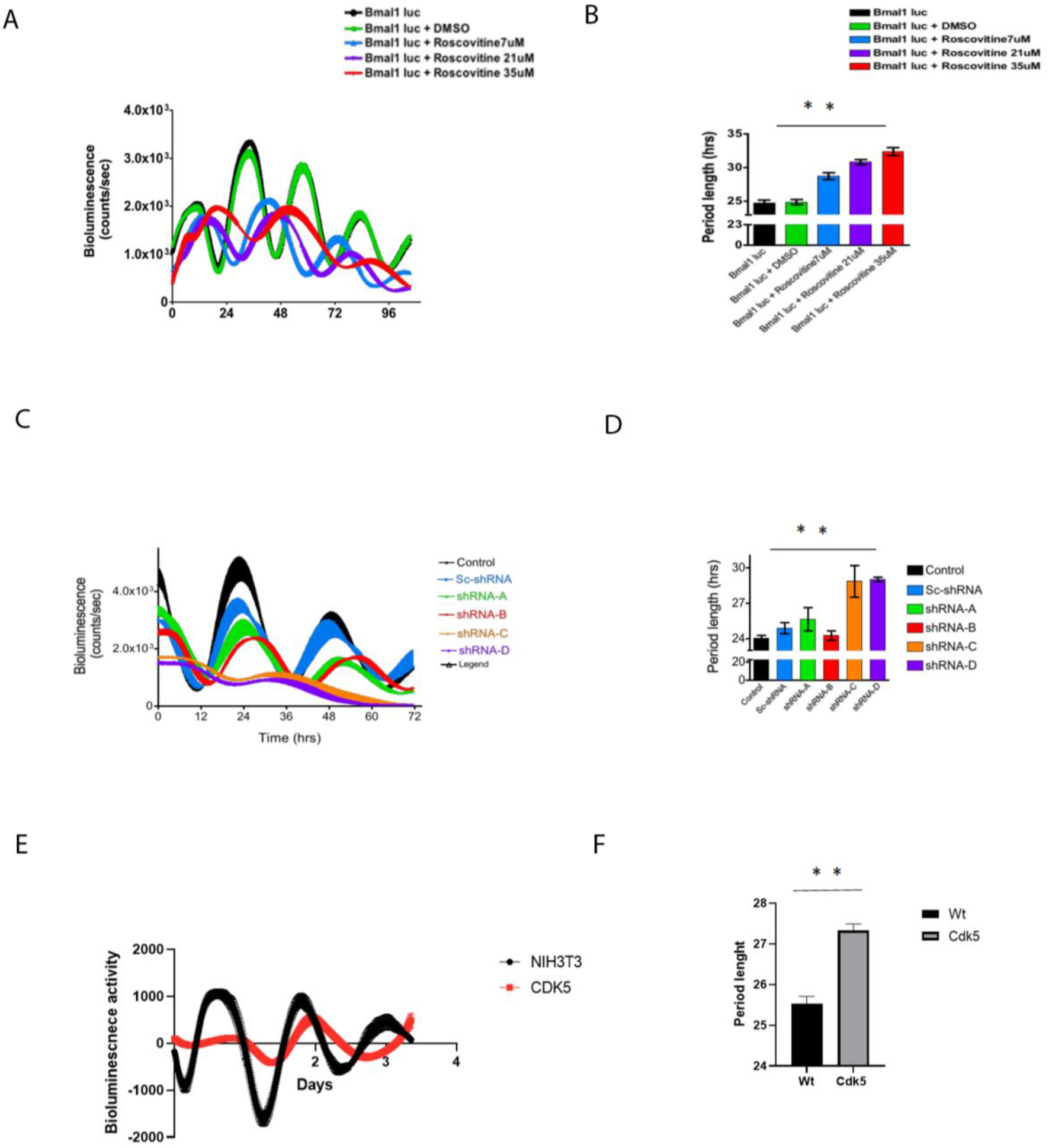
CDK5 affects the circadian clock. A) NIH3T3 cells previously treated with an increasing amount of Roscovitine (7, 21, 34 uM) were transfected with *Bmal* ^*luc*^ promoter and synchronized with forskolin. The bioluminescence was recorded over several days. B) The period length was quantified, and statistical analysis was performed. The results showed that an increased amount of forskolin corresponded to a prolonged period of length. Quantification of three independent experiments (mean ± SEM). One-way ANOVA with Bonferroni’s post-test, *: p<0.001.

### CDK5 affects the clock factors nuclear localization

The lack of CDK5 slows the clock down in the peripheral clocks (Fig.2). Thus, we aimed to define the effect of CDK5 on the clock at the molecular level. So far, only PER2 and CLOCK are directly phosphorylated by CDK5 (Brenna et al., 2019; Kwak et al., 2013). Therefore, we investigated whether both PER2 and CLOCK accumulation and distribution in the NIH 3T3 could be affected in the absence of the kinase. Cells were synchronized with forskolin and collected at different CTs. This samples collection was followed by nucleo/cytoplasm fractionation to observe whether the cellular distribution of CLOCK and PER2 was affected by CDK5. In control cell lines, we could observe PER2 at CT 12,16, CT 28, and 32 in both nuclear and cellular extracts with a peak between CT28-32 (Fig. 3A, B, wt).

**Figure 3.**
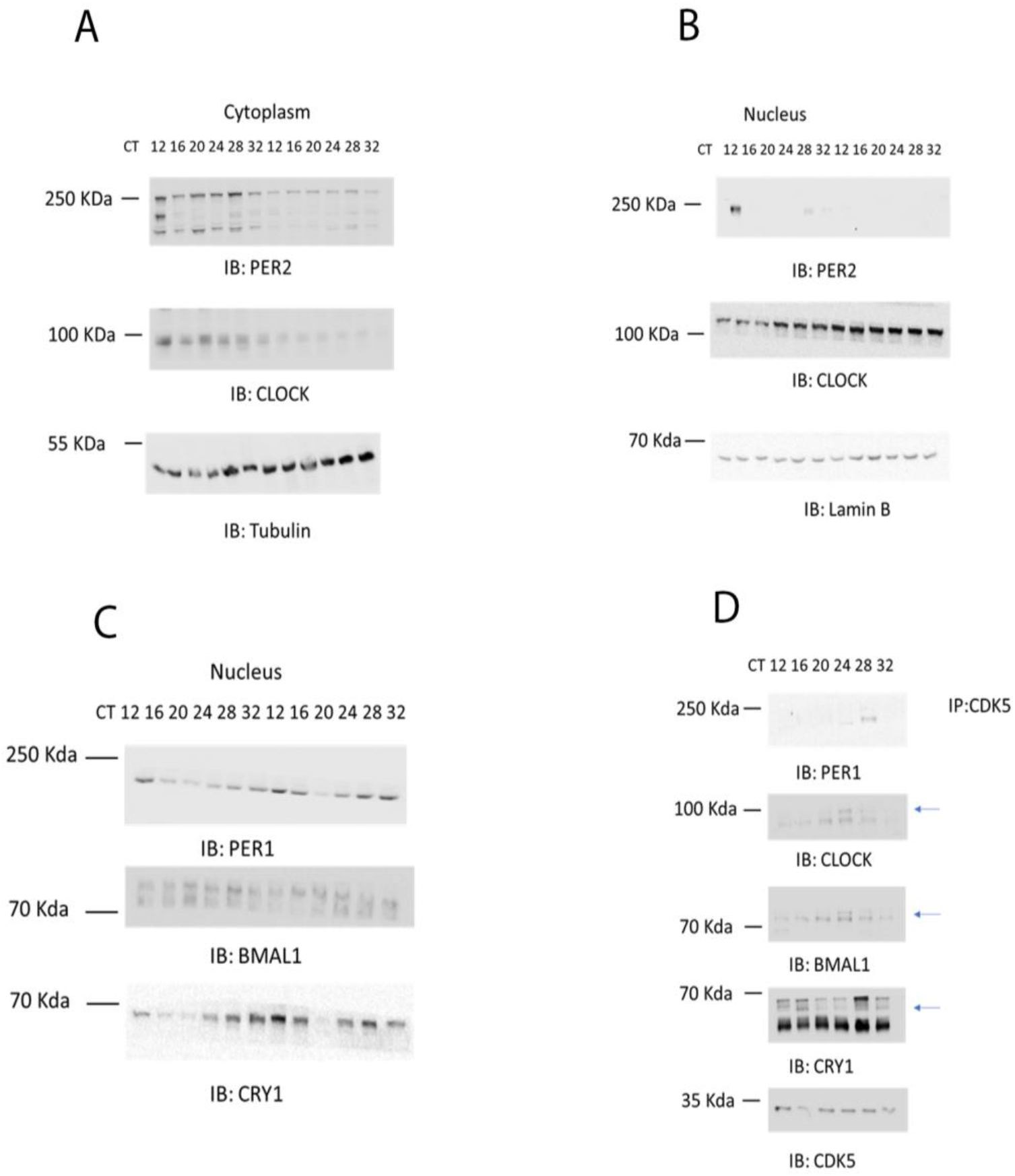
CDK5 affects the clock machinery. A) western blot was performed on cytoplasmic extracts obtained from wt and *Cdk5* ko synchronized cells, showing how the kinase affected the circadian accumulation of CLOCK and PER2. B) western blot performed on nuclear extracts obtained from wt and Cdk5 ko synchronized cells, showing how the kinase affected the circadian accumulation of CLOCK and PER2. C) western blot performed on nuclear extract obtained from wt and Cdk5 ko synchronized cells, showing how the kinase affected the circadian accumulation of BMAL1, CRY1 PER1. D) Immunoprecipitation performed around the clock on wt cells synchronized with forskolin using CDK5 antibody as a bait. CDK5 co-resolved with BMAL1, CLOCK, CRY1, PER1 at specific time points

On the other hand, PER2 was observable, in a reduced amount only at CT 12 in *Cdk5* KO cell lines both in cytoplasmic and nuclear extract (Fig. 3A, B, ko cells). As expected, the protein accumulation did not last longer, confirming that CDK5 is important for PER2 protein stability (Brenna et al., 2019). Subsequently, we analyzed CLOCK cellular distribution. The cytoplasmic protein accumulation showed a flat profile, while rhythmic in the nuclei in wt cell lines peaking between CT 24-32 (Fig. 3A, B wt). By contrast, cytoplasmic accumulation of CLOCK was dampened in *Cdk5* KO cells lines compared to WT while in the nuclei lost the rhythmicity and was way more abundant (Fig. 3A, B, ko cells). Altogether, these results show CDK5 has an opposite effect on CLOCK and PER2 in murine embryonic fibroblasts. Both CLOCK and PER2 are main members of the molecular clock machinery (Arya et al.). Thus, We sought to investigate how the lack of CDK5 could affect the nuclear accumulation of the other proteins part of the machinery, such as BMAL, CRY1, and PER1. Thus, we subsequently investigated whether BMAL, PER1, CRY1 nuclear localization was affected as well. Our immunoblotting showed that BMAL was accumulated in the wt nuclei at CT20, while the protein accumulation was shifted about 4 hours in the *Cdk5* ko cells (Fig. 3C). Furthermore, Immunoblottings using antibodies against PER1 and CRY1 showed that in wt cell lines, the nuclear accumulation of these factors was higher at the subjective night. By contrast, the nuclear accumulation of these proteins in the *Cdk5* ko cell lines was much higher than wt, and we could detect PER1 and CRY1 during the subjective day (FIG. 3D).

Taken together, our results show that CDK5 affect the assembly of the clock machinery in mouse embryonic fibroblast. Due to the severe impact on the clock machinery caused by the lack of CDK5, we wondered whether this enzyme could be part of the clock machinery. There are many examples of kinases being part of the complex in central and peripheral clocks (Brenna et Albrecht 2020). Therefore, we immunoprecipitated CDK5 at different CT, every 4-hours, starting from CT12 to CT32, followed by wb immunodetecting CLOCK, BMAL1, PER1/2, CRY1. Our results showed that CDK5 could interact with PER1, PER2, and CRY1 in the subjective night. CDK5:CRY was observable at CT12-16, while CDK5:PER1 was detected at CT16-20.

Interestingly, both CLOCK and BMAL1 interact with CDK5 during the subjective day, with an identical profile. This observation confirms that CLOCK and BMAL work as a heterodimer (Fig. 3D). Altogether our results show that CDK5 is part of the clock machinery.

### Lack of CDK5 shifts BMAL recruitments to the chromatin and, as a consequence, the mRNA rhythmicity

Our results show that CDK5 is part of the clock machinery, and the lack of this enzyme is responsible for a spatial and temporal redistribution of the complex. We subsequently wondered how this event could affect the expression of the CCGs. To investigate that aspect, we synchronized both wt and *Cdk5* ko cells with forskolin and collected samples every 4 hours starting from CT 12 up to CT40. We subsequently extracted total mRNA and performed RT-qPCR to detect the accumulation of clock factors pre-mRNA. We investigated *Bmal1* gene expression, which is under control of ROR alpha, and, *Per1* which is under the control of CLOCK: BMAL1, and *Clock*, which should not show any oscillation. Our results show interesting molecular behaviors. *Bmal1* gene expression in wt cells peaked around 24 hours after the stimulus. In *Cdk5* ko cells, the peak of pre-mRNA accumulation was shifted 4 hours ahead (Fig. 4A). Interestingly, *Bmal1* expression mirrored the nuclear protein accumulation, suggesting a control mediated by CDK5 on BMAL at the transcriptional level. Furthermore, *Per1* gene expression in both wt and Cdk5 ko cells mirrored exactly the *Bmal1* profile (Fig. 4B). Surprisingly, Clock pre-mRNA, which did not show any oscillation in wt cells, gained ex Novo rhythmicity in the *Cdk5* ko (Fig 4C). Altogether our results show that CDK5 plays a role in the regulation of the clock-controlled genes.

**Figure 4.**
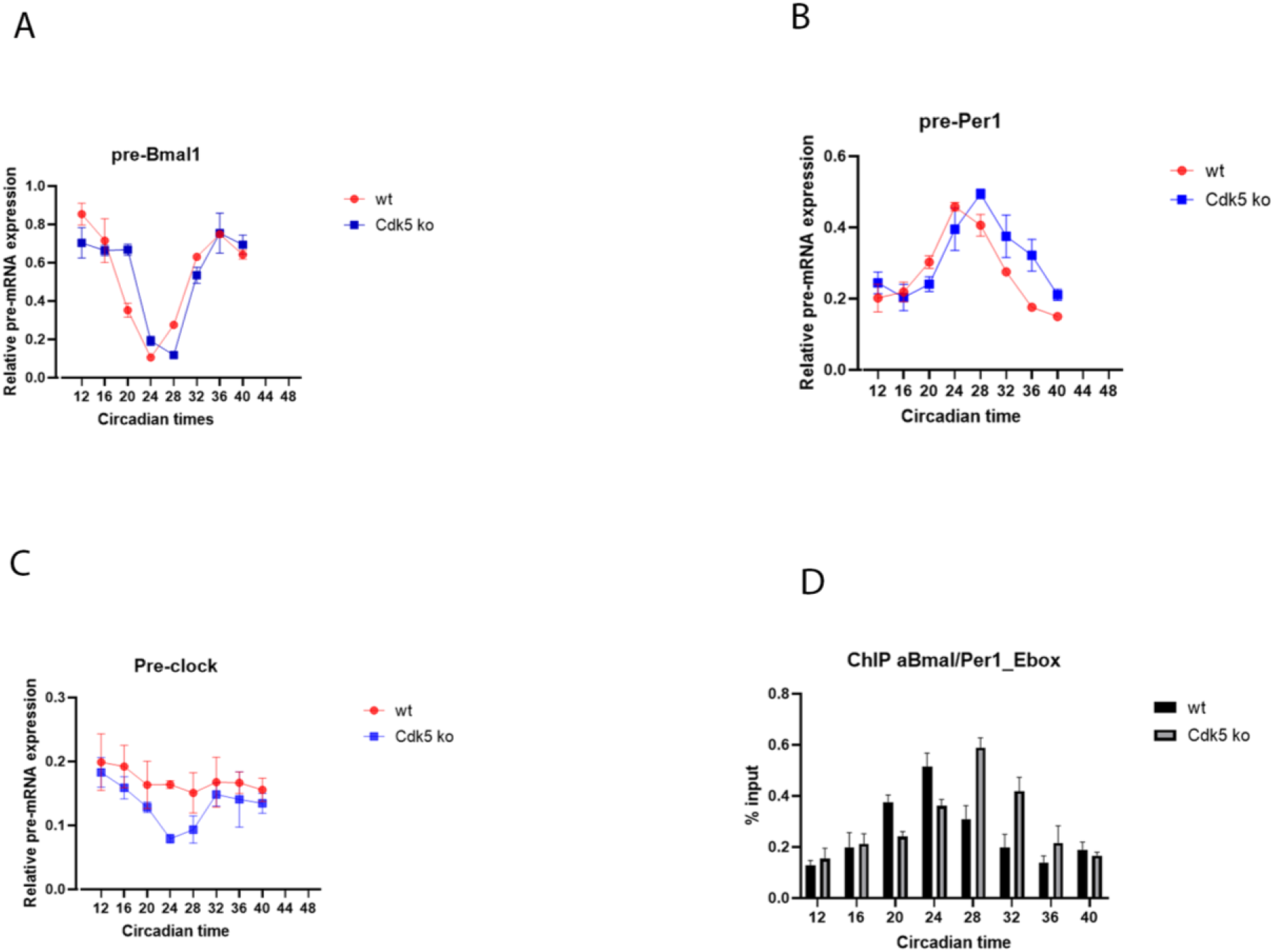
CDK5 influences the expression of clock-controlled genes. The temporal profile of pre-mRNA accumulation for *Bmal1* (A), *Per1* (B), *Clock* (C) was analyzed in synchronized wt and Cdk5 ko cells. Quantification of three independent experiments (mean ± SEM). One-way ANOVA with Bonferroni’s post-test, *: p<0.001 D) Chromatin immunoprecipitation was performed using BMAL1 antibody as a bait. RT-qPCR was performed using specific probes that amplified the E-box of Per1. Quantification of three independent experiments (mean ± SEM). One-way ANOVA with Bonferroni’s post-test, *: p<0.001

Additionally, our previous results showed that CDK5 affects the nuclear localization of the heterodimer CLOC: BMAL1. Particularly, BMAL1 nuclear accumulation in *Cdk5* ko cells is delayed by 4-hours. BMAL1 is the main transcriptional factor promoting the expression of genes whose promoter contains E-boxes (Ripperger and Schibler 2006). Thus, these observations raised whether the different nuclear localization of BMAL would also mirror different temporal recruitment of the transcriptional factor on the E-box of CCGs. We performed chromatin immunoprecipitation (ChiP) to challenge the hypothesis. We immunoprecipitated the transcriptional factor at the same time we analyzed the pre-mRNA profiles before. Thus, we performed RT-qPCR to amplify the E-box within the promoter of the *Per1* gene. Interestingly, our results show that the recruitment of BMAL1 on the *Per1*_E-box was rhythmic in both cell lines. However, while in wt cells, the peak of rhythmicity was at CT24 in the Cdk5 ko, at CT28 (Fig. 4D). Thus, our results showed that CDK5 influences the circadian period in the peripheral via influencing nuclear accumulation and, therefore, temporal promoter occupancy of BMAL1 on the CCGs.

## Discussion

It has been recently described that phosphorylation is the stronger mark of rhythmicity in circadian mammal models (Robles et al., 2017). Furthermore, many kinases were described to influence the circadian clock in livings in the past decades, such as CKI, CDK1, GSK3 B (Brenna et Albrecht 2020).

It was recently showed the impact of Cyclin-Dependent Kinase 5 (CDK5) on the circadian clock in the SCN, which is the peacemaker of circadian rhythmicity (Brenna et al., 2019). Indeed in mice, when CDK5 is silenced in the SCN, it affects PER2 protein accumulation and nuclear translocation. Consequently, mice show a shortening in the period length when their behavior is measured in constant darkness. However, it was described in the recent past that CDK5 could influence many pathways in peripheral tissues, such as regulating vesicular transport, apoptosis, cell adhesion, and migration ((Contreras-Vallejos et al., 2012) Dysregulation of CDK5 can alter these pathways leading to many diseases connected to circadian dysfunctions ((Pozo and Bibb 2016; Sulli et al., 2019).

Since CDK5 has a profound impact on the central clock, we questioned whether it could also affect the clock in the periphery. For this reason, we used mouse embryonic fibroblast cell lines as *in vitro* model system. After synchronization with forskolin, CDK5 shows a higher kinase activity during the subjective night than the subjective day (Fig. 1B, C). Our data align with the diurnal activity of CDK5 in the SCN, which was also peaking at night (Brenna et al., 2019). We further questioned the impact of CDK5 on the circadian rhythmicity in NIH3T3. We performed a Real-Time Bioluminescence measurement using the *Bmal*^*luc*^ promoter as a reporter. We tested the rhythmicity in different conditions: 1) we used Roscovitine which is a potent inhibitor of CDK5 kinase activity (Ref); 2) we used different sh RNA targeting *Cdk5* gene; 3) we used CRISPR/CAS9 Cdk5 KO cell lines.

All the different approaches showed that the lack of CDK5 causes a lengthening of the circadian period in cell lines (Fig 2). This observation came as a surprise since it looks quite the opposite of what was observed before in the SCN of mice (Brenna et al., 2019). This difference shows that CDK5 might have a tissue-specific function. However, a systemic comparison should be performed between the SCN and the MEFs to understand what factor(s) regulated by CDK5 can switch from shorter to longer period length. CLOCK and PER2 are the only described target for CDK5 kinase activity (Brenna et al., 2019; Kwak et al., 2013). Therefore, we performed nuclear/cytoplasm fractionation on synchronized cells to detect the circadian subcellular localization of both CLOCK and PER2 (Fig. 3A, B). We observed less CLOCK in the cytoplasm in the *Cdk5* KO compared to wt.

Moreover, the nuclear accumulation in wt cells showed a circadian profile that overlapped the profile of clock-controlled gene expression such as *Per1* (Fig. 4B). On the other hand, in *Cdk5* ko cells, the profile of protein accumulation was flat, and there was generally more CLOCK at each time point than wt. These results were aligned with previous observations showing that CDK5 influences the nuclear localization of CLOCK (Kwak et al., 2013). These results are interesting since the wider accumulation of CLOCK can be related to the prolonged period length observed in cell culture (Fig.2). Furthermore, since CLOCK is observed as a heterodimer with BMAL1, we also investigated the nuclear accumulation of the transcriptional factor. Our results show that BMAL1 nuclear accumulation in wt cells peaks at CT 20 while Cdk5 ko at CT24, 4-hours later (Fig. 3C).

Interestingly, CRY1 and PER1, members of the repressive loop of the circadian clock, were accumulated in the nuclei of the wt cells with a coherent profile, peaking at the subjective night. However, in the ko cell lines, these factors were highly accumulated also during the subjective day (Fig. 3C). Since CDK5 affected so strongly the cellular localization of all the clock factors described above, we wondered whether the kinase per se could be part of the clock machinery. Therefore we performed an immunoprecipitation around the clock to define the temporal profile of interaction between CDK5 and the other clock factors. Surprisingly, CDK5 interacts with CRY1 at the early night, while with PER1 at the late night. CDK5 also interacts with both CLOCK and BMAL simultaneously during the subjective day, confirming that CLOCK and BMAL1 work as heterodimers (Wang et al., 2013). Thus, CDK5 seems to be part of the clock machinery. Other kinases are part of the complex, such as Casein Kinase I (Aryal et al., 2017), which directly phosphorylates both PER1 and PER2 (Camacho et al., 2001).

As a consequence, the expression of the clock-controlled genes is also affected. For example, *Bmal1* gene expression shows a 4-hours delayed peak in *Cdk5* ko cells. Since *Bmal1* gene expression is regulated by the stabilizing loop formed by Rora and Reverb alpha (Akashi et al., 2005), this result suggests that CDK5 might be part also of the stabilizing complex. *Per1* expression also shows 4-hours of delay in the peak in Cdk5 ko cells, which mirrored the profile of BMAL1 nuclear accumulation. Thus, together we show that the shift of BMAL1 accumulation in Cdk5 Ko cells overlaps the shift in the expression of target genes. To finally confirm that the shift in the gene expression of *Per1* is related to delayed recruitment of BMAL1 onto its E-box in Cdk5 ko cells, we performed a ChIP assay using an antibody against BMAL1. Our ChIP assay showed that while BMAL1 is recruited onto the E-box in wt cells with a circadian profile peaking at CT24, the peak was observed at CT28 in the ko cell lines. Thus, The recruitment of BMAL was shifted by 4 hours (Fig. 4D). Finally, we acquired another interesting observation. Clock gene expression, which showed a flat profile in wt cells, gained an ex Novo oscillation in the *Cdk5* ko cells. Altogether, our evidence shows that CDK5 plays an important role in the peripheral clock using the cell culture model system. However, it is not clear how the lack of this enzyme can provoke a shortening of the period length in the SCN, while the opposite in the cells. It would also be interesting how this mechanism of regulation described in this paper would influence specific organs such as the liver, gut, muscles, testis, which show all independent oscillations (Dibner et al., 2010)

## MATERIALS and METHODS

### Cell culture

Mouse embryonic fibroblasts (MEFs) were obtained from mice without functional CDK5 using the CRISPR/CAS9 approach. The details are explained in Brenna et al., 2019. All lines were maintained in Dulbecco’s modified Eagle’s medium (DMEM) containing 4.5 g/l glucose,stable glutamine, sodium pyruvate, and 3.7 g/l NaHCO_3_ (PAN Biotech), supplemented with 10% fetal calf serum (FCS) and 100 U/mL penicillin-streptomycin at 37°C in a humidified atmosphere containing 5% CO_2_. Forskolin stimulation (100 μM) was used to synchronized the cellular circadian rhythm. Samples were collected at specific time points mentioned in the text.

### *In vitro* kinase assay immunoprecipitating CDK5 from cells

CDK5 was immunoprecipitated from cells samples at different circadian times (CTs) (circa 200 μg of protein extracts. After immunoprecipitation at 4°C overnight with 2x Protein A agarose (Sigma-Aldrich), samples were diluted in washing buffer and split in two halves. One half of the IP was used for an in vitro kinase assay. Briefly, 1 μg of histone H1 (Sigma-Aldrich) was added to the immunoprecipitated CDK5 and assays were carried out in reaction buffer (30 mM Hepes, pH 7.2, 10 mM MgCl_2_, and 1 mM DTT) containing [γ-^32^P] ATP (10 Ci) at room temperature for 1 hr and then terminated by adding SDS sample buffer and boiling for 5 min. Samples were subjected to 15% SDS-PAGE, stained by Coomassie Brilliant Blue, and dried, and then phosphorylated histone H1 was detected by autoradiography. The other half of the IP was used for Western blotting to determine the total amount of CDK5 immunoprecipitated from the SCN samples. To quantify the kinase activity at each time point, the following formula was used: ([^32^P] H1/total H1 for each reaction)/CDK5 IP protein.

### RNA extraction from cells

Cells were grown to confluency on 6 cm Petri dishes and induced with 10 μM forskolin (50 mM stock in dimethyl sulfoxide) for the indicated time. Total RNA was extracted using the Nucleospin RNA II kit (Macherey & Nagel) and adjusted to 1 μg/ μl with water. An amount of 1 μg was reverse-transcribed using Superscript II with random hexamer primers (Thermo Fisher). Real-time PCR was performed using the KAPA probe fast universal master mix and the indicated primers on a Rotorgene 6000 machine. The relative expression was calculated compared to the geometric mean of expression of the inert genes Nono, SirT2, Atp5h, and Gsk3b ^69^. For a complete list of primers used in the paper, please see Table 1.

### Chromatin Immunoprecipitation

Chromatin immunoprecipitation from cells was performed as described before ^70^. Briefly, the cells were grown to confluency on 15 cm Petri dishes, induced with 10 μM forskolin (50 mM stock in dimethyl sulfoxide), and fixed at the indicated time with 1% formaldehyde/ 1x phosphate-buffered saline buffer (PBS) for 10 min at 37°C. Then the cells were washed twice with ice-cold 1x PBS and scraped in 1 ml of 10 mM dithiothreitol/ 100 mM tris(hydroxymethyl)aminomethane (Tris) HCl, pH 8.8. Cells were lysed in 10 mM ethylenediaminetetraacetic acid (EDTA), 1 mM ethylene glycol-bis(2-aminoethyl ether)-N, N, N’, N’-tetraacetic acid (EGTA), 10 mM 4-(2-hydroxyethyl)-1-piperazineethanesulfonic acid (HEPES), pH 7.6, 0.2% Triton X-100 for 5 min on ice and the obtained nuclei washed with 1 mM EDTA, 0.5 mM EGTA, 200 mM NaCl, 10 mM HEPES, pH 7.6. The purified nuclei were sonicated in 2 mM EDTA, 150 mM NaCl, 20 mM Tris, pH 7.5, 1% SDS using a BRANSON SLPe sonicator with a 4C15 tip for 6 cycles of 10 s each kept on ice, and diluted 1:10 with 2 mM EDTA, 150 mM NaCl, 20 mM Tris, pH 7.5, 1.1% Triton X-100. Equal amounts of chromatin were incubated with the indicated antibodies for 1h at RT. (Table 2) and the immune complexes were captured with protein A agarose fast-flow beads (Sigma-Aldrich) for 1h at RT. The beads were washed with 2 mM EDTA, 150 mM, 20 mM Tris, pH 7.5, 0.1% SDS, 1% Triton X-100, then 2 mM EDTA, 500 mM, 20 mM Tris, pH 7.5, 0.1% SDS, 1% Triton X-100, then 2 mM, 250 mM LiCl, 20 mM Tris, pH 7.5, 0.5% Na-deoxycholate, 0.5% NP40 substitute, and finally 2 mM EDTA, 150 mM, 20 mM Tris, pH 7.5. DNA fragments were eluted, and the crosslinks reversed in 2 mM EDTA, 150 mM NaCl, 20 mM Tris, pH 7.5, 1% SDS at 65°C overnight. The DNA fragments were purified using a MinElute PCR fragment purification kit (Qiagen). Real-time PCR reactions were performed using the KAPA probe fast universal master mix and the indicated primers on a Rotorgene 6’000 machine. The efficiency of the precipitations was calculated by comparing the amount of precipitated material to 1% of the starting material.

### Protein extraction from cells

Total confluent cells plated in 10 cm dishes were washed two times with 1x PBS (137 mM NaCl, 7.97 mM Na_2_HPO_4_ × 12 H_2_O, 2.68 mM KCl, 1.47 mM KH_2_PO_4_). Then PBS was added again, and plates were kept for 5 min at 37°C. Cells were detached and collected in tubes and frozen in liquid N_2_. They were subsequently resuspended in Ripa buffer (50 mM Tris-HCl pH7.4, 1% NP-40, 0.5% Na-deoxycholate, 0.1% SDS, 150 mM NaCl, 2 mM EDTA, 50 mM NaF) with freshly added protease or phosphatase inhibitors, and homogenized by using a pellet pestle. Homogenates were kept on ice for 15 min followed by sonication (10 s, 15% amplitude). Right after, the samples were centrifuged for 15 min at 16,100 g at 4°C. The supernatant was collected in new tubes and pellet discarded.

### Immunoprecipitation

Circa 400 μg of protein extract was diluted with the appropriate protein lysis buffer in a final volume of 250 μl and immunoprecipitated using the indicated antibody (ratio 1:50) at 4°C overnight on a rotary shaker. The day after, samples were captured with 50 μl of 50% (w/v) protein-A agarose beads (Roche) for 3 hr at 4°C on a rotary shaker. Before use, beads were washed three times with the appropriate protein buffer and resuspended in the same buffer (50% w/v). The beads were collected by centrifugation and washed three times with NP-40 buffer (100 mM Tris-HCl pH7.5, 150 mM NaCl, 2 mM EDTA, 0.1% NP-40). After the final wash, beads were resuspended in 2% SDS 10%, glycerol, 63 mM Trish-HCL pH 6.8, and proteins were eluted for 15 min at RT. Laemmli buffer was finally added, samples were boiled 5 min at 95° C and stored at -20° C.

### Nuclear/Cytoplasm fractionation (Brenna et al., 2019)

Tissues or cells were resuspended in 100 mM Tris-HCl pH 8.8/10 mM DTT and homogenized with a disposable pellet pestle. After 10 min incubation on ice, the samples were centrifuged at 2500 g for 2 min at 4°C and the supernatant discarded. After adding 90 μL of completed cytoplasmic lysis buffer (10 mM EDTA, 1 mM EGTA, 10 mM Hepes pH 6.8, 0.2% Triton X-100, protease inhibitor cocktail (Roche), NaF, PMSF, ß-glycerophosphate), the pellet was resuspended by vortexing, followed by centrifugation at 5200 rpm for 2 min at 4°C. The supernatant transferred into a fresh 1.5 mL tube was the CYTOPLASMIC EXTRACT. The pellet was washed three times with cytoplasmic lysis buffer and resuspended in 45 μL 1x NDB (20% glycerol, 20 mM Hepes pH 7.6, 0.2 mM EDTA, 2 mM DTT) containing 2x proteinase and phosphatase inhibitors followed by adding 1 vol of 2x NUN (2 M Urea, 600 mM NaCl, 2% NP-40, 50 mM Hepes pH 7.6). After vortexing the samples were incubated 30 min on ice, centrifuged 30 min at 13,000 rpm at 4°C and the supernatant that was transferred into a fresh tube was the NUCLEAR EXTRACT.

### Western blot

The indicated amount of protein was loaded onto 10% SDS-PAGE gel and run at 100 Volt for two hours. Once the migration was completed, we performed a semidry transfer (40 mA, 1 hour 30 s) using Hybond® ECL™ nitrocellulose membranes followed by red ponceau staining (0,1 % of Ponceau S dye and 5% acetic acid) to validate the success of the transfer. The membrane was subsequently washed with TBS 1x/Tween 0.1% and blocked with TBS 1x/Milk 5%/Tween 0.1% for 1 hour. After washing, the membrane was stained with the appropriate primary antibodies overnight. The antibodies against all the clock factors were homemade (final dilution 1:1000). CDK5 purchased by Cell Signaling Tech (1:1000), while Tubulin by Abcam (1:1000) and Lamin B by Santa Cruz (1:1000). The day after, membranes were washed three times with TBS 1x/Tween 0.1% followed by secondary antibody immunoblotting for 1 hour at room temperature. The densitometric signal was digitally acquired with the Azure Biosystem.

### Statistical analysis

Statistical analysis of all experiments was performed using GraphPad Prism8 software. Depending on the type of data, either an unpaired t-test with Welch’s correction or a paired t-test, when a sample was set to 1, were performed. Two-way ANOVA was performed on time and genotype-dependent experiments. Values considered significantly different are highlighted, *p<0.05, **p<0.01, or ***p<0.001.

## Acknowledgments

Support by the Swiss National Science Foundation (310030_184667/1) and the State of Fribourg is gratefully acknowledged.

## Author Contributions

Conceived and designed the experiments: JR, AB.

Performed the experiments: JR, AB.

Analyzed the data:, JR, AB.

Contributed reagents, materials, analysis tools: UA

Wrote the paper: AB

Edited the manuscript: AB

